# Host targets of effectors of the lettuce downy mildew *Bremia lactucae* from cDNA-based yeast two-hybrid screening

**DOI:** 10.1101/863738

**Authors:** Alexandra J.E. Pelgrom, Claudia-Nicole Meisrimler, Joyce Elberse, Thijs Koorman, Mike Boxem, Guido Van den Ackerveken

## Abstract

Plant pathogenic bacteria, fungi and oomycetes secrete effector proteins to manipulate host cell processes to establish a successful infection. Over the last decade the genomes and transcriptomes of many agriculturally important plant pathogens have been sequenced and vast candidate effector repertoires were identified using bioinformatic analyses. Elucidating the contribution of individual effectors to pathogenicity is the next major hurdle. To advance our understanding of the molecular mechanisms underlying lettuce susceptibility to the downy mildew *Bremia lactucae,* we mapped a network of physical interactions between *B. lactucae* effectors and lettuce target proteins. Using a lettuce cDNA library-based yeast-two-hybrid system, 61 protein-protein interactions were identified, involving 21 *B. lactucae* effectors and 46 unique lettuce proteins. The top ten targets based on the number of independent colonies identified in the Y2H and two targets that belong to gene families involved in plant immunity, were further characterized. We determined the subcellular localization of the fluorescently tagged target proteins and their interacting effectors. Importantly, relocalization of effectors or targets to the nucleus was observed for four effector-target pairs upon their co-expression, supporting their interaction *in planta*.

## Introduction

Plant pathogenic bacteria, fungi and oomycetes deploy effector proteins to manipulate host cell processes. Importantly, effectors serve to suppress and circumvent plant immune responses. Basal host defense responses are activated upon recognition of ubiquitously present microbe-associated molecular patterns (MAMPs) by plant pattern recognition receptors (PRRs). Evolutionary adapted pathogens deploy effectors to suppress this pattern-triggered immunity (PTI). Specialized intracellular nucleotide-binding and leucine-rich repeat receptors (NLRs) recognize host translocated effectors, or the perturbations effectors induce on host proteins, resulting in the activation of effector-triggered immunity (ETI). In turn, ETI can be counteracted by other effectors leading to a state of effector-triggered susceptibility (ETS) [1].

Fungi and oomycetes secrete apoplastic effectors that operate at the host-pathogen interface, and host-translocated effectors that act intracellularly in the host. The genomes of different pathogens encode for extensive candidate effector sets, which have specific characteristics based on their origin. Fungal genomes encode e.g. small apoplastic cysteine-rich proteins [2], plant pathogenic downy mildews and *Phytophthora* species express host-translocated Crinklers and RXLR effectors [3–6] whereas plant pathogenic Gram-negative bacteria, e.g. *Pseudomonas syringae,* inject type III effectors into host cells [7]. At present, the major challenge lies in elucidating the contribution of individual effectors to the infection process through the identification of plant targets and analysis of the molecular mechanisms that contribute to disease susceptibility.

Effector targets were systematically studied in *Arabidopsis thaliana* by identifying physical interactions between *Arabidopsis* proteins and effector proteins of the bacterium *P. syringae*, the obligate biotrophic oomycete *Hyaloperonospora arabidopsidis*, and the obligate biotrophic ascomycete *Golovinomyces orontii* resulting from a yeast-two-hybrid (Y2H) screening of ∼8000 Arabidopsis ORFs [8, 9]. Interactions between 123 effectors and 178 Arabidopsis proteins were found. Nine Arabidopsis proteins interacted with effectors from all three pathogens, whereas another 24 proteins interacted with effectors from two of the three pathogens. Arabidopsis proteins that interacted with effectors from multiple pathogens were proposed to function as cellular hubs on which effectors of different microbial kingdoms converge to effectively undermine plant immune responses [8]. Disease assays with one or more pathogens were performed on Arabidopsis insertion mutants corresponding to 124 targets and an altered susceptibility phenotype was observed in mutant lines for 63 targets. Susceptibility phenotypes were more frequently observed in mutant lines corresponding to targets that interacted with multiple effectors [9].

The obligate biotrophic oomycete *Bremia lactucae,* the causal agent of downy mildew disease of lettuce, is a major problem in lettuce production worldwide. Consequently, a multitude of genetic studies have been carried out resulting in the identification of over 50 genes mediating resistance to *B. lactucae* in lettuce [10]. On the pathogen side, research efforts have led to the identification and cloning of one Crinkler and 49 *B. lactucae* RXLR-like effectors in *B. lactucae* isolate Bl:24 [11, 12]. To gain insight into the molecular mechanisms underlying lettuce susceptibility to *B. lactucae* infection, we here describe the identification of interactions between 21 *B. lactucae* effectors and 46 unique lettuce proteins, uncovered using the Y2H system. The subcellular localization of twelve targets, selected on Y2H performance and predicted biological function, as well as their interacting effectors was visualized by confocal fluorescence microscopy. Upon co-expression of these effectors and their lettuce targets in *Nicotiana benthamiana*, relocalization of the effector or target to the nucleus was observed in four instances, providing additional evidence that these interactions occur *in planta*.

## Materials & methods

### Generation of the Y2H prey library and bait constructs

A *Lactuca sativa* cv. Olof cDNA library was constructed using Invitrogen Custom Services (Invitrogen, Carlsbad, CA). Briefly, RNA was isolated using phenol/chloroform extraction from mock-treated seedlings, *Bremia lactucae* isolate Bl:24 (compatible interaction) and isolate F703 (incompatible interaction)-infected seedlings at 3 days after inoculation (dpi), and seedlings treated with the salicylic acid (SA) analog benzothiadiazole (BTH; 0.1 mg/ml) 24h after spraying. RNA originating from the different treatments, was mixed in equal amounts. A three-frame uncut oligo(dT)-primed cDNA library in pENTR222 was created from 2 mg RNA using Gateway cloning technology. The library was transferred into the yeast-two-hybrid destination vector pDEST22 to generate GAL4 activation domain (AD) lettuce fusion proteins. The pDEST22 library in *E. coli* strain DH10B originated from 22 x 10^6^ colony forming units (cfu) with an average insert size of 1.1 kb.

*B. lactucae* effectors were amplified using cDNA from the sequence encoding the predicted signal peptide cleavage sites and new start codons were introduced. Gateway entry clones of *B. lactucae* effectors were recombined with the pDEST32 yeast-two-hybrid destination vector using LR clonase to generate GAL4 DNA binding domain (DBD) effector fusion proteins.

### Yeast strains and transformation

To create competent yeast, cells were grown o/n in 250 ml YEPD at 28 °C and 200 rpm to an OD_600_ of 0.2-0.8. Cells were spun down at 1800 rpm for 5 min, washed with 50 ml sterile ddH_2_O, spun down again and washed with 50 ml TE/LiAc (100 mM LiAc, 10 mM Tris, 1 mM EDTA, pH 8.0). After a final centrifugation step, the yeast was resuspended in TE/LiAc to an OD_600_ of 50. For single construct transformation, 20 μl of competent yeast was gently mixed with 11 μl 10xTE (100 mM Tris, 10 mM EDTA, pH 8.0), 13 μl 1 M LiAc, 82 μl 60% PEG (MW 3,350), 20 μl salmon sperm DNA (Sigma #D1626; 2 mg/ml in TE, heated at 95 °C for 5 min and transferred to ice) and 200 ng plasmid. Reactions were incubated at 30 °C for 30 min, and then transferred to a water bath at 42 °C for 15 min. 1 ml ddH_2_O was added to each transformation reaction, the tubes were spun down at 5000 rpm for 30 sec and the pellet was resuspended in ddH_2_O. Bait strains were plated on synthetic complete (Sc) –Leu medium and prey strains were plated on Sc –Trp medium. Colonies appeared after two to three days at 30 °C. For library transformation the protocol was scaled up to 3200 μl competent yeast cells and 90 μg plasmid DNA. Yeast colonies were harvested in YEPD medium + 20% glycerol and 1 ml aliquots with OD_600_ = 40 were frozen at −80 °C. The yeast prey library consisted of 1.1 x 10^6^ – 1.5 x 10^6^ individual colonies. Yeast strain Y8800 (genotype MATa trp1−901 leu2−3,112 ura3−52 his3− 200 gal4Δ gal80Δ cyh2R GAL1::HIS3@LYS2 GAL2::ADE2 GAL7::LacZ@met2) was used for prey and yeast strain Y8930 (genotype MATα trp1−901 leu2−3,112 ura3−52 his3−200 gal4Δ gal80Δ cyh2R GAL1::HIS3@LYS2 GAL2::ADE2 GAL7::LacZ@met2) was used for bait. Bait strains that grew on Sc –Leu –His plates in the absence of prey were considered auto-activating and discarded.

### Library screening

The mating method was used for library screening [13]. Diploid yeast cells were plated on Sc –Leu – Trp –His medium to identify interacting bait-prey combinations. Retransformation of bait and prey candidates was performed to confirm interactions. A detailed description can be found in S1 file.

### Bioinformatic analysis of effectors and lettuce targets

Lettuce prey gene models were extracted using a BlastN search against the lettuce genome [14]. Where necessary, incomplete gene models were corrected using the lettuce transcriptome and prey sequencing data. Prey sequences with a stop codon within the first 50 aa after the GAL4-AD sequence were considered not in frame. Presence of signal peptides was predicted using SignalP4.1 [15] and transmembrane domains were identified with TMHMM [16, 17]. Transmembrane domains identified in effector sequences were validated using TOPCONS [18, 19]. Domain prediction was performed using InterProScan5. The presence of importin-α dependent nuclear localization signals was predicted using cNLS Mapper [20]. WY domains were predicted using a HMM model [21] with an individual cut-off E-value of 0.001 for the best motif within an effector. Post-translational lipid modifications were predicted using GPS-Lipid/CSS-Palm [22, 23] on effector sequences lacking the signal peptide.

*L. sativa* cv. Salinas coding sequences corresponding to genome v8 (Genome ID 28333) were downloaded from the CoGe platform (https://genomevolution.org/). Translated CDSs of targets on the shortlist were searched using Hidden Markov Models for the presence of domains with *E*-value 1e^-4^ as the cut-off. In general, all gene models with the same Pfam domain were subsequently extracted to identify the corresponding gene families. All gene models belonging to the same gene family as the targets on the shortlist were named collectively. Gene names are composed of the prefix ‘*Ls*’ for *L. sativa* followed by a short abbreviation denoting the domain and a number. Genes were numbered according to their position on the lettuce linkage groups 1-9 to avoid issues due to the, often, complex orthologous relationship with Arabidopsis proteins. An overview of all the new nomenclature for gene models can be found in S2 File. A detailed description of the procedure per gene model can be found in S1 File.

### Transient expression in *N. benthamiana*

Full-length prey sequences were amplified from lettuce cv. Olof cDNA or prey plasmid using primers listed in S1 Table and recombined in a modified pGemTEasy vector containing the pDONR201 Gateway recombination site (pGemTEasy^mod^) using BP clonase. Bait and prey entry clones were further recombined in pUBN-YFP-DEST and pUBN-CFP-DEST (kind gift from dr. Christopher Grefen; [24]) or with pB7WGY2 and pB7WGC2 [25] to create N-terminal YFP and CFP fusion proteins respectively using LR clonase and transformed in *Agrobacterium tumefaciens* strain C58C1 (pGV2260) with selection on rifampicin (50 µg/ml), carbenicillin (50 µg/ml) and spectinomycin (100 µg/ml). Leaves of four to five-week-old *N. benthamiana* plants were co-infiltrated with an *A. tumefaciens* strain carrying the P19 silencing suppressor [26] in combination with strains harbouring bait or prey fusion constructs resuspended in infiltration buffer (10 mM MES, 10 mM MgCl2, and 150 µM acetosyringone, pH 5.6) to an OD of 0.3 per *A. tumefaciens* strain.

### Confocal fluorescence microscopy

Microscopy was performed at 2-3 days after *Agrobacterium*-infiltration using a Zeiss LSM 700 laser scanning microscope. Leaf sections were incubated in propidium iodide (PI) solution (5 mg/ml) for 7-10 min prior to imaging to stain the cell wall. Excitation of Cyan Fluorescent Protein (CFP), Yellow Fluorescent Protein (YFP) and PI was done at 405 nm, 488 nm and 555 nm respectively. Emitted light of CFP and YFP was captured using a 490-555 nm band-pass filter, whereas emitted light of PI was captured using a 560 nm long pass filter.

### Immunoblotting

Leaves were harvested 2-3 days after infiltration with *Agrobacterium* suspensions, frozen in liquid nitrogen and stored until further processing at −80°C. All subsequent steps were performed at 4°C. Leaves were ground using mortar and pestle and proteins were extracted with lysis buffer (10 mM Tris-HCl pH 7.6, 150 mM NaCl, 0.5 mM EDTA, 0.25% (v/v) Triton-X 100, 0.25% (v/v) Tween-20, 5 mM DTT, 4% (w/v) polyvinylpyrrolidone) supplemented with 2x protease inhibitor cocktail without EDTA (#11873580001, Roche) for 1 h at 4°C in an Eppendorf ThermoMixer at 2000 rpm. Lysates were centrifuged for 15 min at 10 000 g. The supernatant was separated on Bolt pre-mixed cast gradient gels 4-12% with Bolt Mes-SDS running buffer (ThermoFisher Scientific) according to the manufacturer’s instructions. Proteins were transferred for 100 min at 100 V to nitrocellulose membrane using Towbin buffer (25 mM Tris, 192 mM glycine, 20% (v/v) ethanol), detected with 1:2500 diluted anti-GFP HRP conjugated antibodies (Miltenyi Biotec Cat# 130-091-833, RRID:AB_247003, Lot no: 5180709500) using SuperSignal West Pico chemiluminescent substrate (ThermoFisher Scientific) and documented with a CCD-camera (ChemiDoc, Bio-Rad).

## Results

### Effector interaction screening identifies new and known target candidates

To investigate the molecular mechanisms underlying lettuce susceptibility to *B. lactucae* infection, we selected 46 previously identified candidate effectors from *B. lactucae* isolate Bl:24 [11,12,27] to be used as bait in Y2H screens (Table 1). Orthologs of 34 effectors were found in isolate SF5 [28]. Consistent with the biotrophic lifestyle of *B. lactucae*, the vast majority of these candidate effectors are classified as RXLR-like effectors; when we started the Y2H screen only a single Crinkler (*<u>B</u>remia <u>l</u>actucae* <u>C</u>rinkler) had been discovered. The RXLR-like effectors can be subdivided in two groups: 1) effectors with an RXLR motif (BLR effectors) or RXLR-like motif such as GXLR (BLG effectors) and QXLR (BLQ effectors), and 2) effectors that show homology to RXLR effectors but lack an RXLR-like motif, the BLN (*<u>B</u>remia <u>l</u>actucae* <u>N</u>o RXLR motif) effectors. All of the BLN effectors and the majority of the effectors with an RXLR(-like) motif contain an EER(-like) motif. The effectors ranged in size from 65 to 814 amino acids with an average length of 248 aa.

**Table 1.**
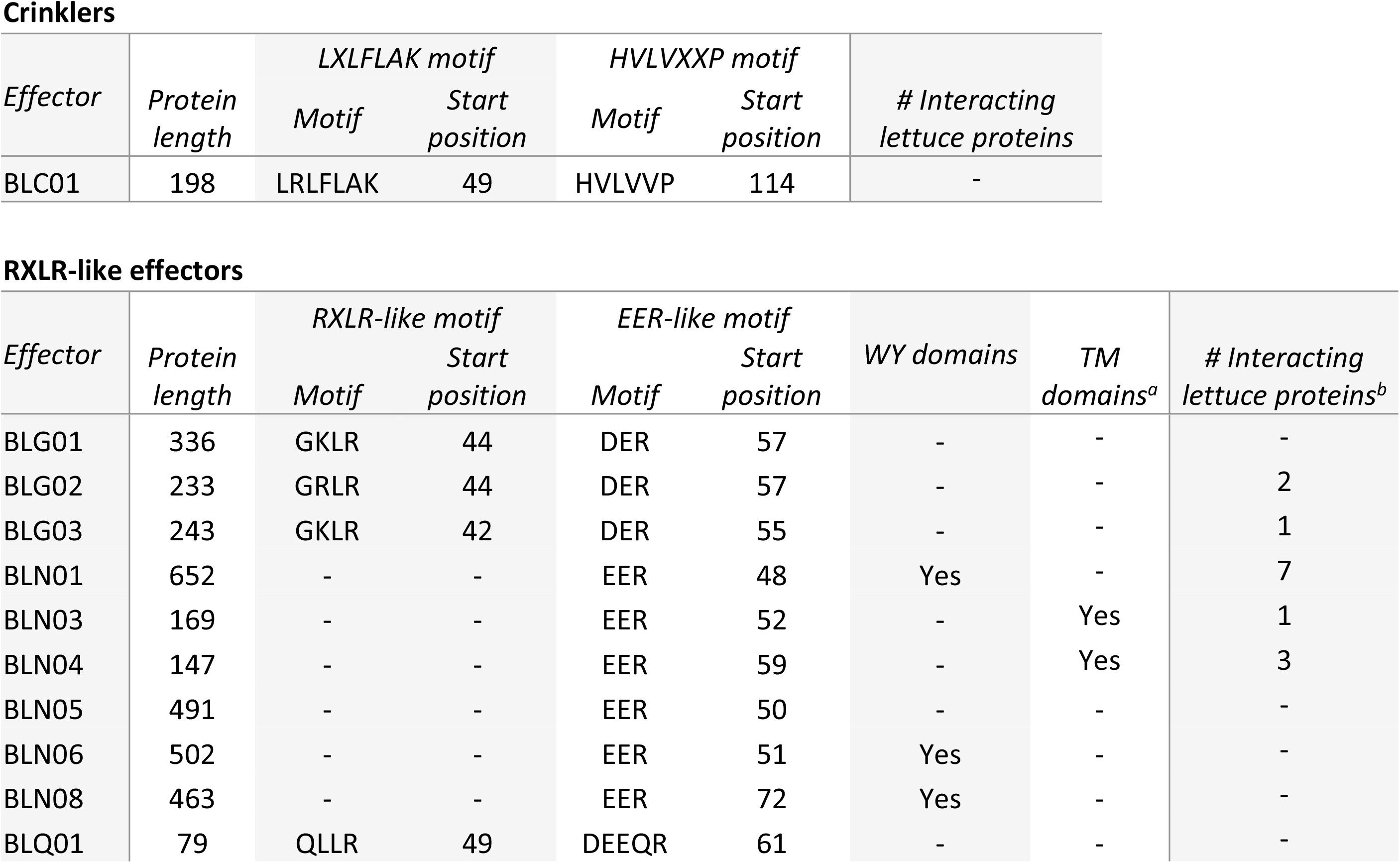
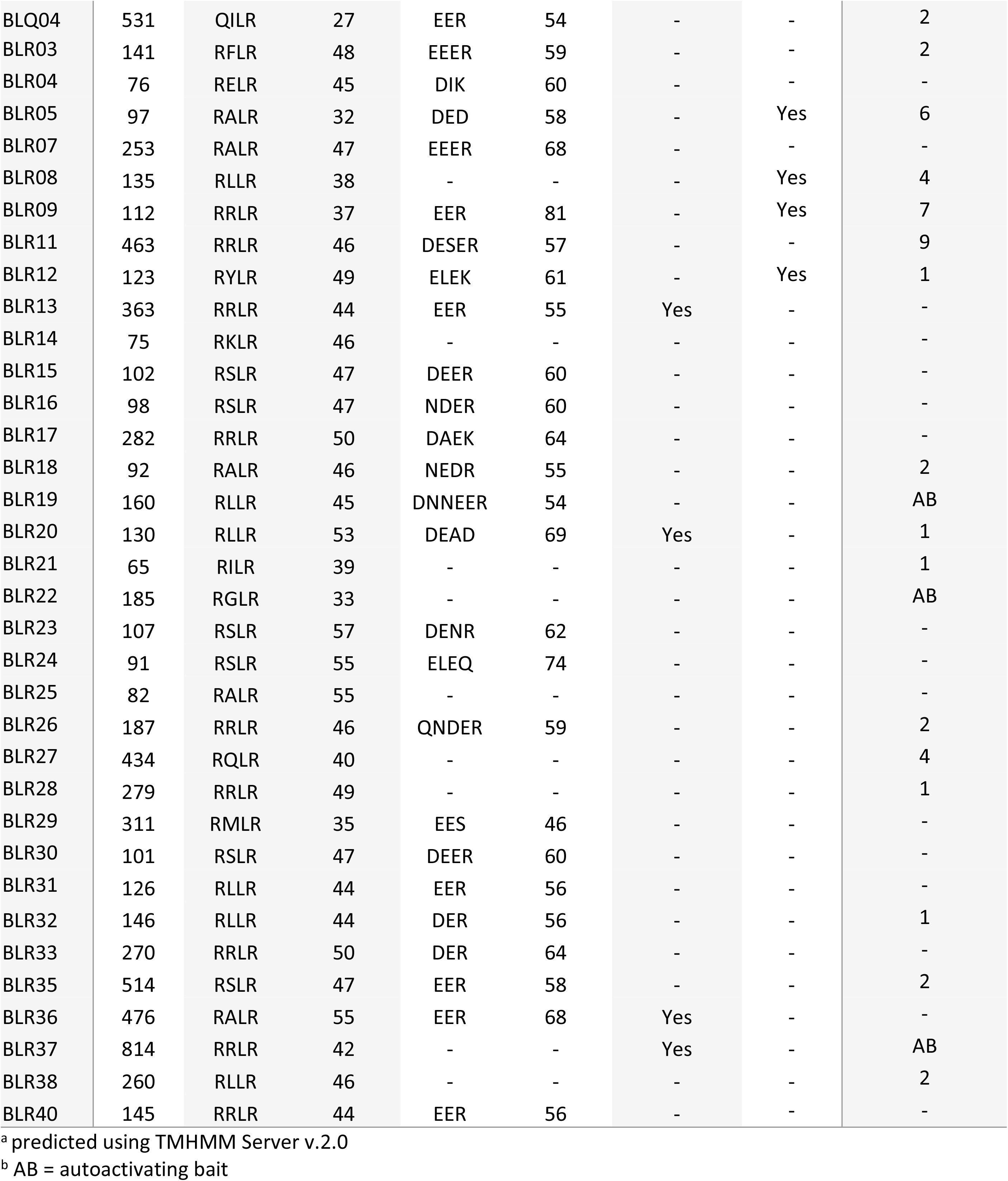
Overview of Bremia lactucae effectors and the number of interacting plant proteins by Y2H screening.

Despite a lack of general sequence conservation between all effector proteins within a species, it was found that about 44% of the *P. infestans* effectors, 26% of *H. arabidopsidis* effectors and 18% of *Plasmopara halstedii* RxLR effectors contain WY-domains that form structurally conserved α-helical folds [21,29,30]. Analysis of the 46 *B. lactucae* effectors revealed WY domains in 7 effectors (15%) (Table 1). Among the WY effectors are BLN06 and BLN08 that lack an RXLR motif, but are nevertheless recognized in specific lettuce lines [12, 27], thereby demonstrating effector activity.

To identify lettuce proteins that are targeted by *B. lactucae* effectors a Y2H screen was performed. First, the coding sequences of the 46 *B. lactucae* effectors lacking the signal peptide-encoding part were cloned in frame with the GAL4-DBD in the bait vector pDEST32. Of the 46 bait constructs three showed auto-activation in the Y2H system. The remaining 43 effectors were used as baits in Y2H screens with a prey library composed of *L. sativa* cv. Olof complementary DNA (cDNA). To promote representation of lettuce transcripts relevant to host-pathogen interactions, lettuce seedlings were challenged with compatible or incompatible *B. lactucae* isolates or treated with a SA-analog before RNA extraction. To eliminate false positives, identified candidate targets from lettuce were carefully scrutinized. Firstly, coding sequences that were out of frame with the GAL4 activation domain-encoding sequence were discarded. Secondly, prey constructs that tested positive for autoactivation of the *HIS3* reporter in the absence of the corresponding bait construct were also removed. Thirdly, bait and prey plasmids were retransformed in yeast to test if the interaction could be confirmed. These selection steps resulted, predominantly, in the elimination of candidate effector targets that were identified only once. In contrast, most candidate effector targets identified in two or more yeast colonies proved reliable interactors. Therefore, all candidate targets that were identified only once were omitted from further analysis. Ultimately, the library screens and subsequent validation steps resulted in interactors for 21 effectors (46%) (Table 1) providing a set of 46 unique interacting lettuce proteins (Table 2).

**Table 2.**
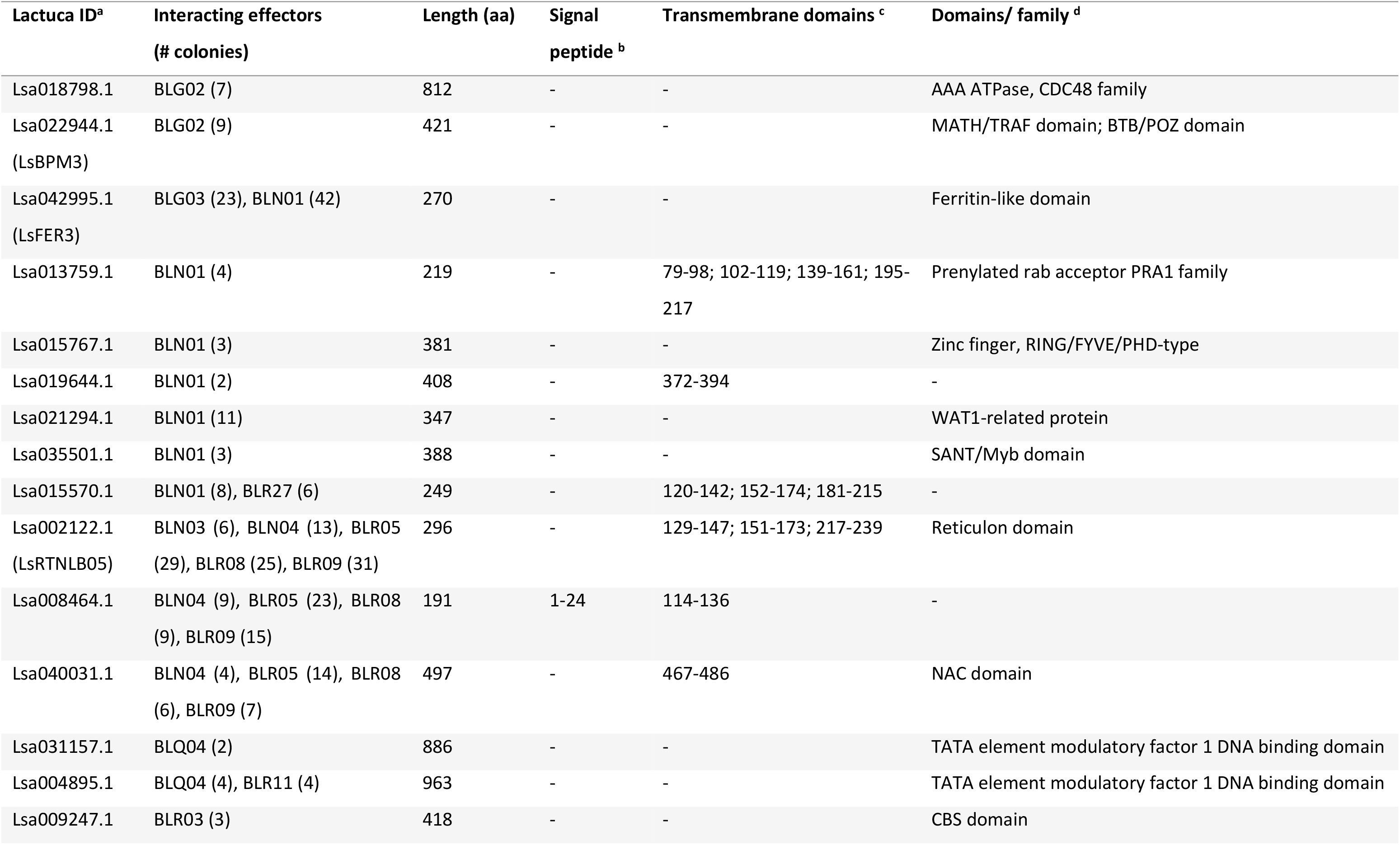
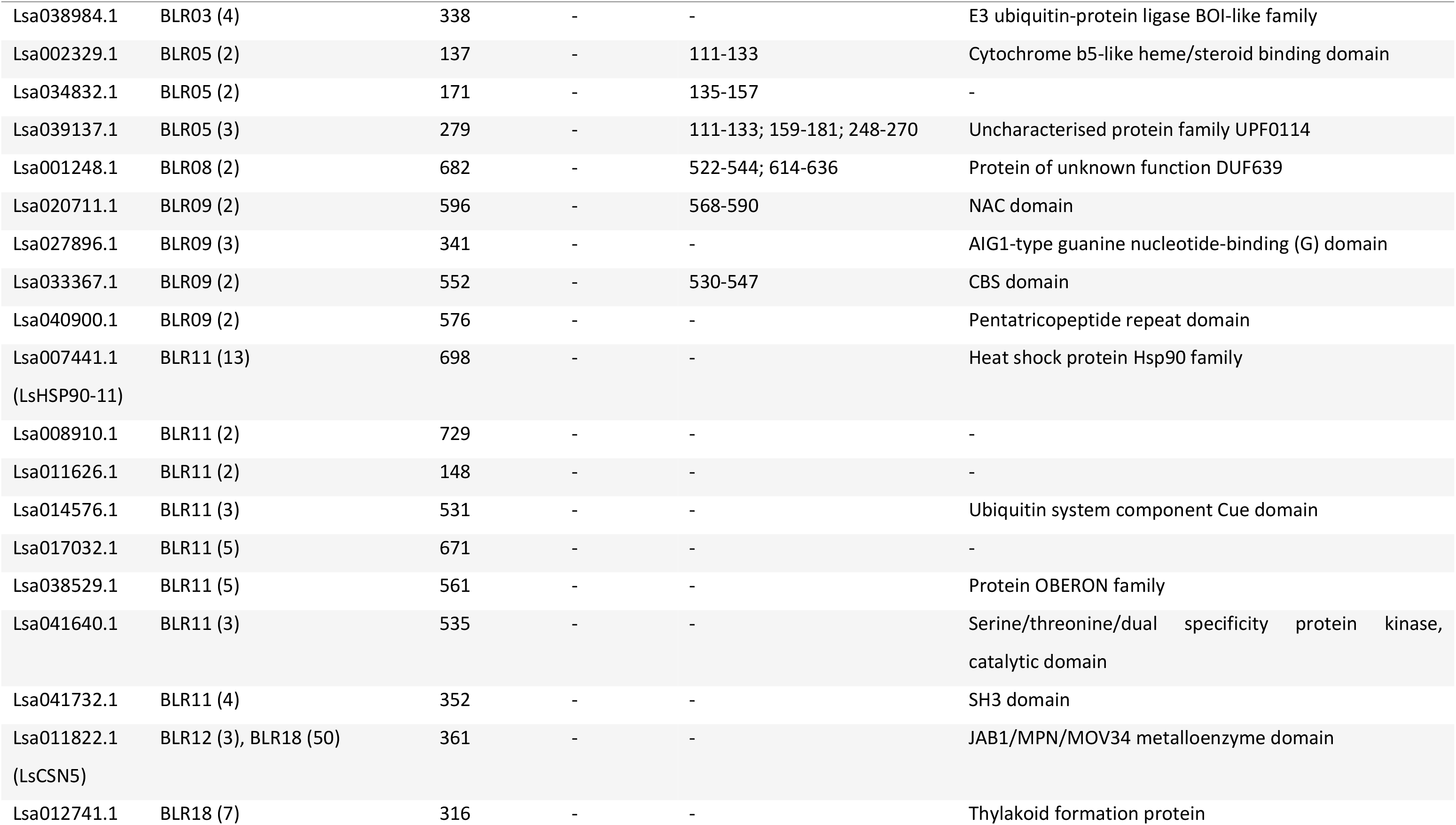
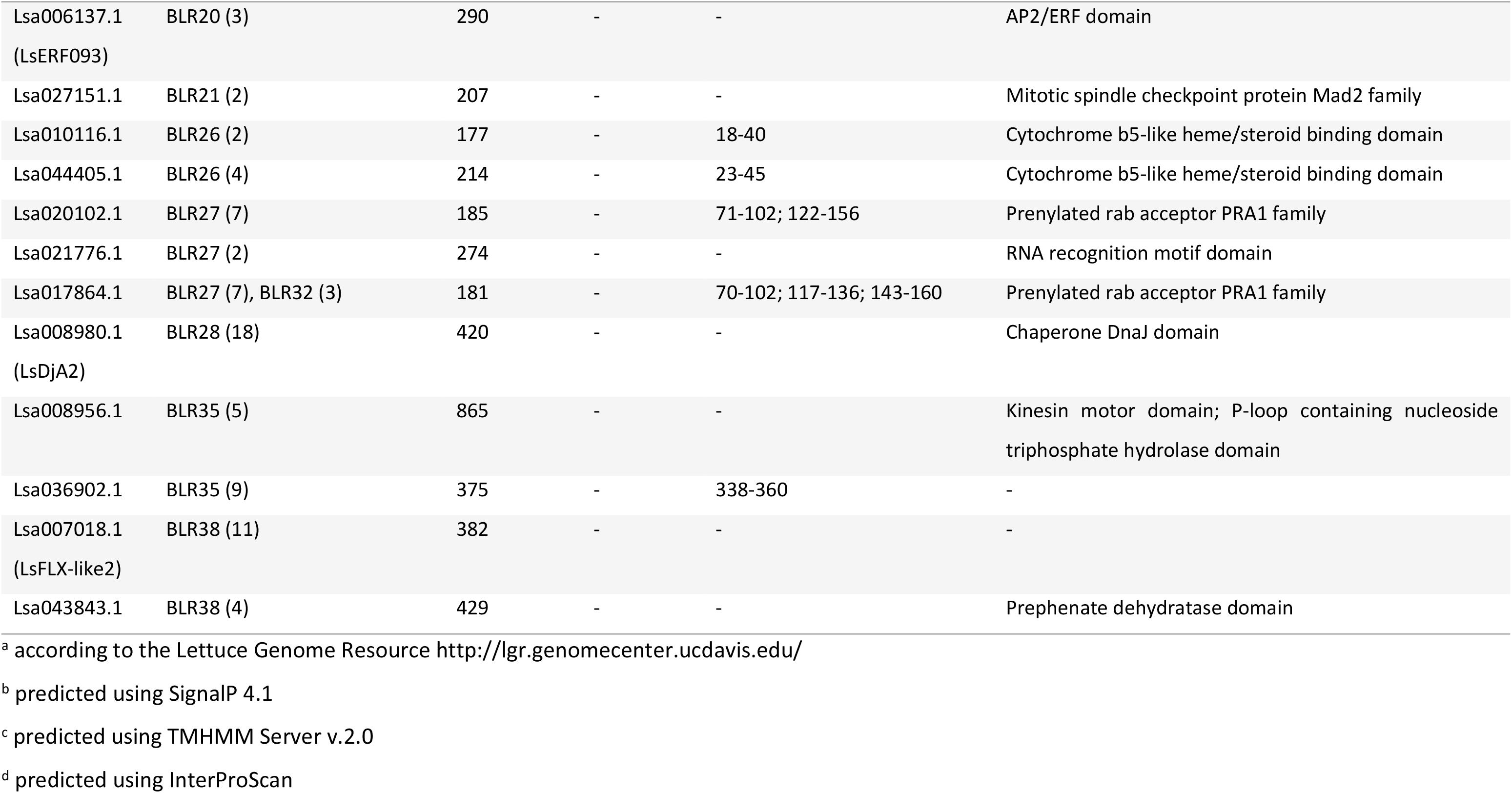
Lettuce proteins identified in yeast-two-hybrid screens.

Interactions between effectors and lettuce proteins were further classified according to the number of unique colonies corresponding to the same lettuce gene and the interaction strength based on reporter gene activation (Fig 1). Due to the stringent selection criteria, only two weak (activation of *HIS3* reporter only) interactions (3%) were identified, 25 interactions (41%) were classified as intermediate (activation of *HIS3* reporter in the presence of 2 mM 3AT) and 34 interactions (56%) were strong (activation of both the *HIS3* and *ADE2* reporters). The vast majority of interactions between effectors and lettuce targets were highly specific: only eight preys (17%) interacted with multiple effectors. LsRTNLB05, a reticulon-like protein, stood out from other lettuce targets due to its interaction with five effectors and a combined total of 104 yeast colonies (Table 2). The most yeast colonies, 50, within a single effector screen were identified with effector BLR18 for LsCSN5 that encodes COP9 signalosome subunit 5 (CSN5). However, LsCSN5 also displayed weak activation of the *HIS3* reporter in the presence of empty bait vector.

**Fig 1.**
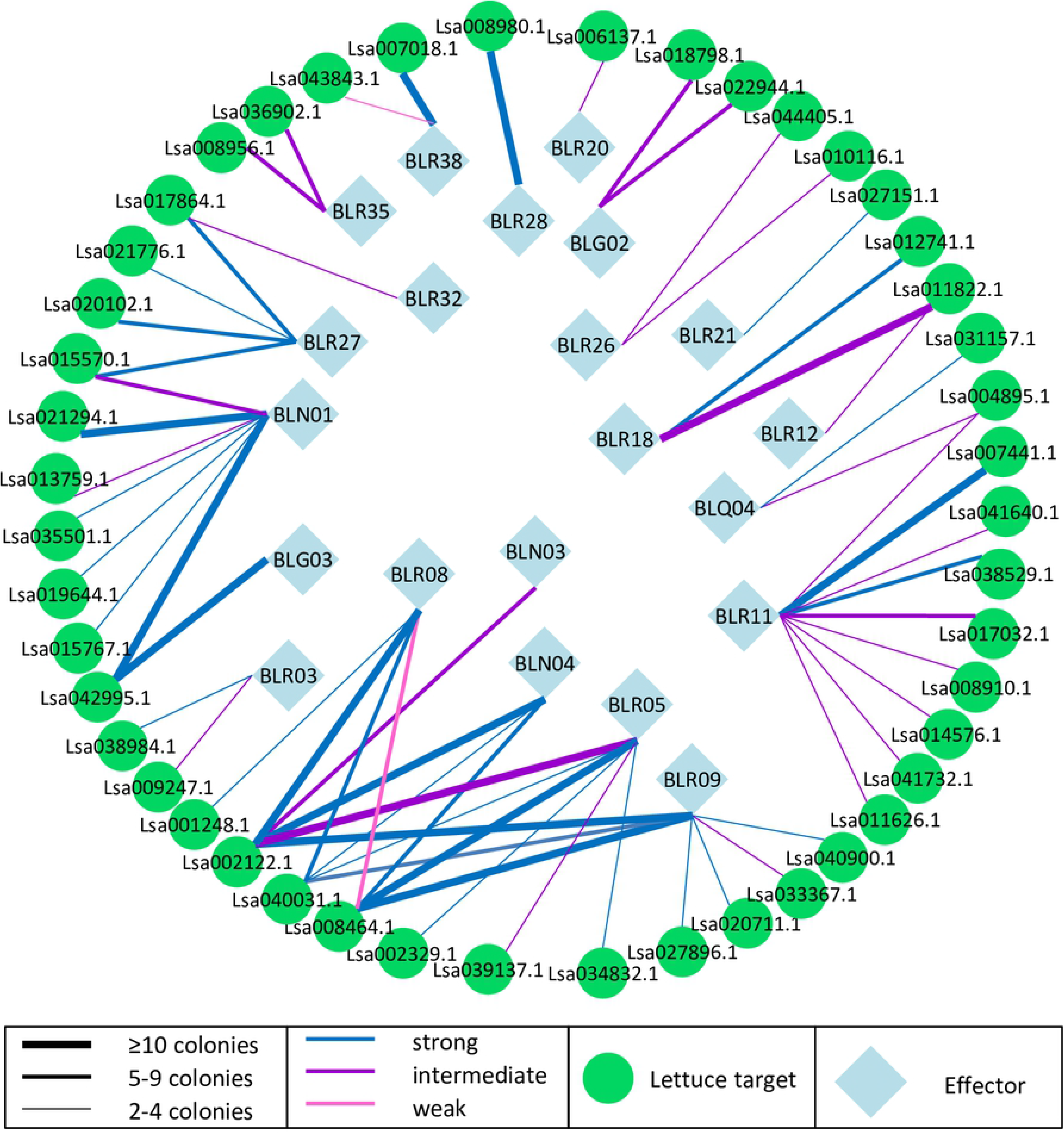
Yeast-two-hybrid identified *B. lactucae* effector – lettuce protein network. Interactions between *B. lactucae* effectors (blue diamonds) and lettuce proteins (green circles) are depicted according to the number of independent colonies representing a lettuce target and the level of reporter gene activation.

The most represented family of proteins in the screen is the prenylated rab acceptor (PRA1) family for which fragments of six members were identified and three members passed all selection criteria. *PRA1* genes encode small transmembrane proteins that localize to the secretory pathway in Arabidopsis and are proposed to play a role in vesicular trafficking in plants [31]. Also, they have previously been described as targets of *G. orontii* effector candidates in Arabidopsis [9]. In lettuce the PRA1 protein family is composed of 17 members. The six Y2H identified PRA1 proteins were found in four clades of the phylogenetic tree (Fig 2A). To determine the specificity of the interactions between *B. lactucae* effectors and PRA1 proteins, a targeted Y2H assay was performed. All six identified prey constructs containing lettuce *PRA1* gene fragments were cotransformed with three interacting and one non-interacting *B. lactucae* effector in yeast and plated on selective medium of increasing stringency. The negative control BLR09 did not interact with any of the PRA1 members. The interactions between effectors BLR27, BLR32 and BLN01 with individual PRA1 proteins as identified in the library screens were confirmed. Furthermore, BLR27 and BLR32 interacted with more PRA1 proteins than determined by the library screen, indicating that the Y2H screening was not exhaustive and not all possible interactions were captured. In contrast, BLN01 interacted robustly with LsPRA1.B1 but only weakly with other PRA1 proteins (Fig 2B). Two of the three PRA1 proteins that passed all selection criteria belong to clade F. Interestingly, the Arabidopsis PRA1 proteins that interacted with *G. orontii* and *H. arabidopsidis* effectors also belong to clade F [9].

**Fig 2.**
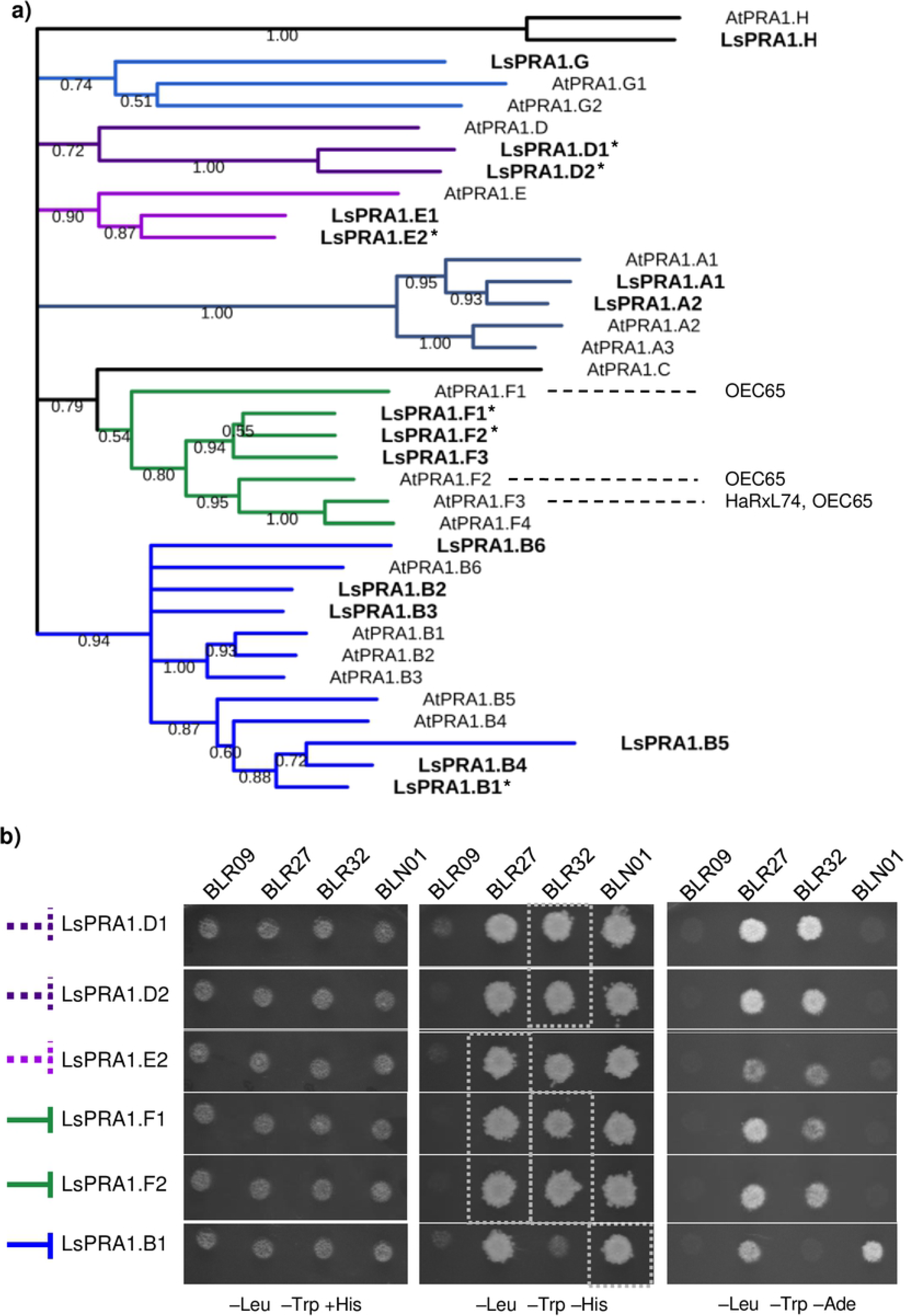
Three *B. lactucae* effectors interact with multiple PRA1 family members. A) Phylogenetic clustering of 17 lettuce and 19 Arabidopsis PRA1 domain-containing proteins. Sequences were aligned using Clustal Omega. Tree construction was performed in MEGA 7.0 using Neighbor-Joining with partial deletion of sites with less than 95% coverage. Nodes with less than 50% confidence score based on 1000 bootstrap replicates were removed. The previously reported interactions between *Hyaloperonospora arabidopsidis* and *Golovinomyces orontii* effectors HaRxL74 and OEC65 and Arabidopsis PRA1 proteins have been added. An asterisk (*) behind the protein indicates it was included in the targeted Y2H. B) A targeted Y2H was performed with yeast isolated prey plasmids containing gene fragments of six PRA1 family members using *B. lactucae* effectors BLR27, BLR32 and BLN01 as bait. BLR09 was included as a negative control bait. Dashed line indicates the PRA1 protein did not pass all Y2H selection criteria, a continuous line indicates the PRA1 proteins passed the Y2H selection criteria. Left: permissive plate containing histidine. Middle: moderately selective plate lacking histidine. Right: strongly selective plate lacking adenine. The positions of previously identified interactions in the library screens are indicated with grey blocks for selection on SC medium lacking histidine.

Twelve plant targets were selected for validation of protein-protein interactions *in planta*, including the top ten ranked effector targets by total number of yeast colonies. In addition, LsBPM3 and LsERF093 were chosen because of their predicted functions in targeting proteins for degradation and transcriptional regulation respectively, as will be described in more detail later. Using domain prediction and orthologous relationships to Arabidopsis proteins, nine out of the twelve were given gene names.

To further explore the selected effector targets, a closer look was taken at the prey plasmid-encoded gene fragments. DNA sequences obtained by Sanger sequencing were aligned to the corresponding lettuce coding sequences. Lettuce gene fragments that included the start codon of the lettuce coding sequence (Fig 3) were detected for eight preys. Yeast clones of the remaining four preys only contained fragments that started downstream of the predicted start codon. The use of a cDNA library that contains both full-length coding sequences and gene fragments likely increased the number of identified protein-protein interactions.

**Fig 3.**
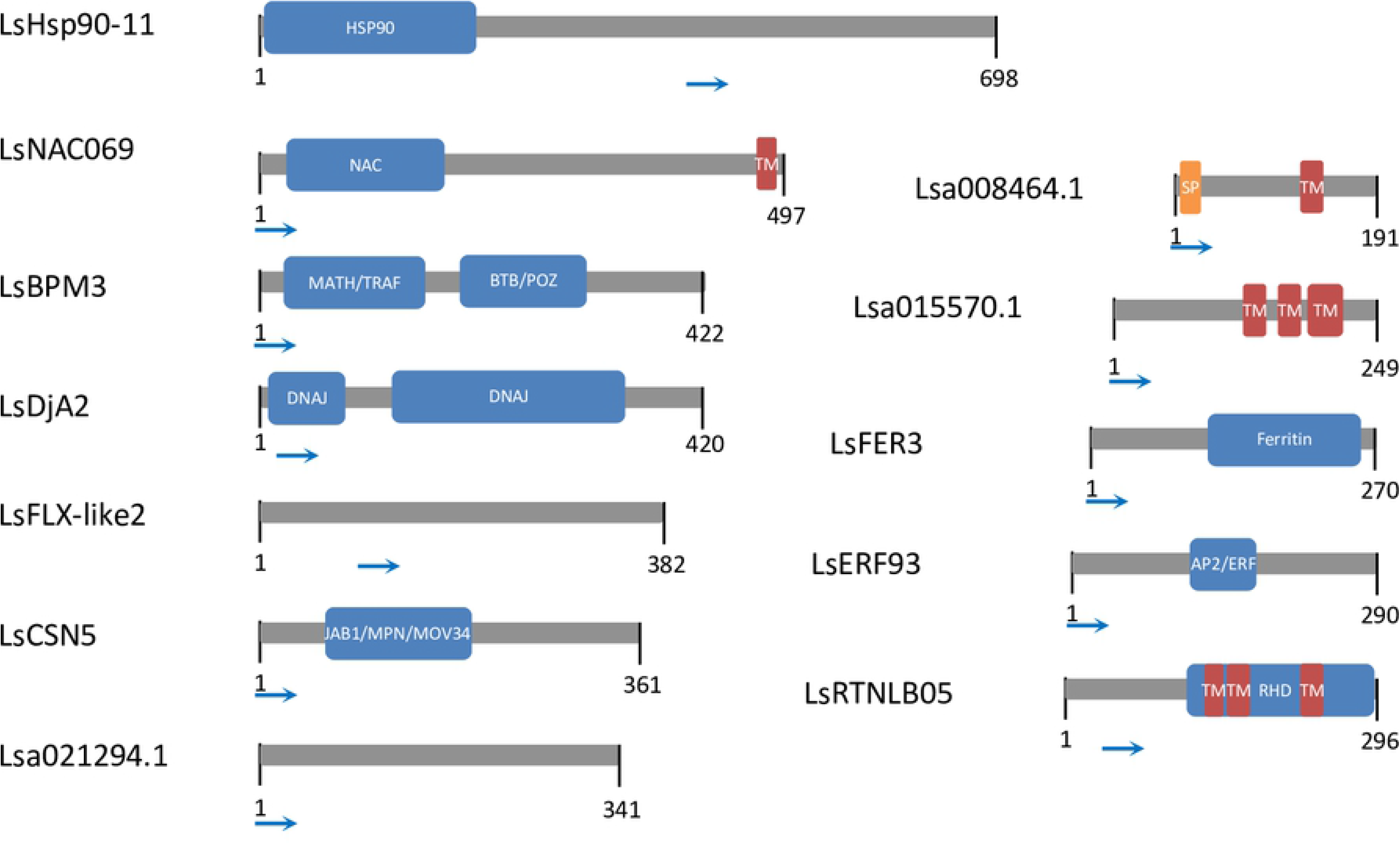
Graphical representation of Y2H-identified effector targets. The start position of the longest identified cDNA fragment is indicated with a blue arrow for each ORF. Domains were predicted using InterProScan. TM = transmembrane domain, SP = signal peptide

### Colocalization of effectors and their plant targets

As proteins that are localized in the same subcellular compartment *in planta* are more likely to be true interactors, the subcellular localization of *B. lactucae* effectors and their plant targets was determined using confocal fluorescence microscopy. The full-length coding sequences of the selected lettuce genes were cloned and fused downstream of *CFP*. Effector sequences were fused to *YFP*. The fusion proteins were produced in leaves of *Nicotiana benthamiana* using *Agrobacterium*-mediated transient expression. Immunoblots were performed to determine the size and stability of the fusion proteins.

Of the 15 effectors described here, four localized exclusively to the cytoplasm and two showed both cytoplasmic and nuclear localization (Fig 4 and Table 3). Immunoblots of YFP-BLR18 showed a large fraction of free YFP and a minor fraction corresponding to the intact fusion protein. Therefore, it is possible that the observed nucleocytoplasmic signal does not reflect YFP-BLR18 but mainly free YFP. YFP-BLR28 and BLR38-YFP contain predicted nuclear localization signals (NLSs) (S2 Table) and localized solely to the nucleus (also see [12]). Six effectors contain a predicted C-terminal transmembrane domain (Table 1 and S3 Table) and were thus expected to localize to membranes. We were unable to detect BLN03 and BLN04 C- or N-terminal fusion proteins. YFP-BLR05 and YFP-BLR09 predominantly labeled the endoplasmic reticulum (Fig S1), whereas YFP-BLR08 was associated with ring-like structures of various sizes in line with our previously reported findings [32]. YFP-BLR12 appeared as punctate structures. Upon prolonged overexpression, YFP-BLR12 also labeled the endoplasmic reticulum suggesting that the punctate structures represent components of the secretory pathway. However, this requires confirmation using appropriate markers. Interestingly, BLG03 that is recognized in *Dm2* lettuce lines [11], does not contain predicted transmembrane domains, but YFP-BLG03 was associated with the plasma membrane (Fig S1) similarly to BLN06-YFP and BLR40-YFP [12]. It is possible that post-translational modifications (S4 Table) contribute to observed membrane localizations.

**Fig 4.**
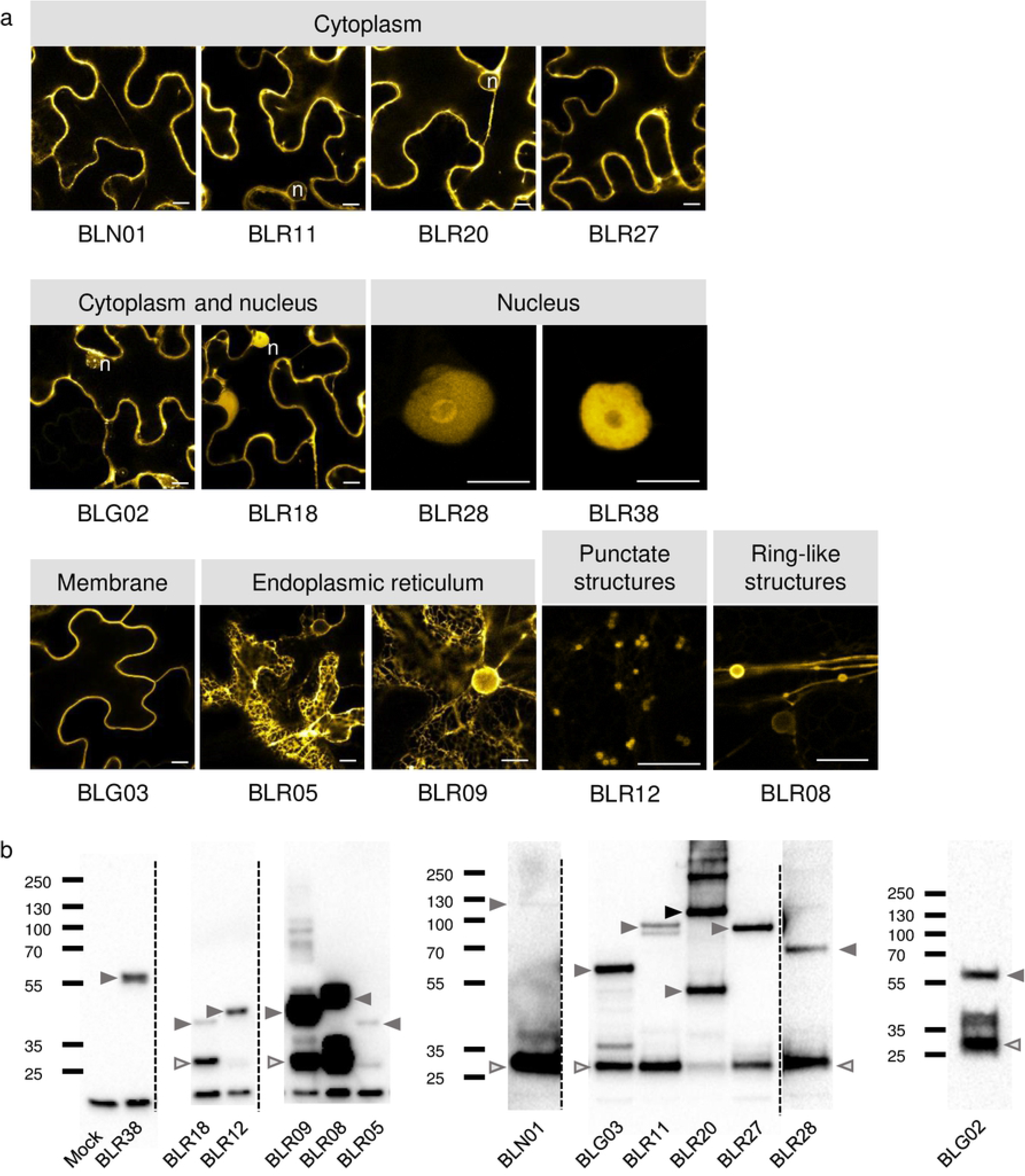
Subcellular localization and stability of *B. lactucae* effectors in *N. benthamiana*. Effectors were fused N-terminally to YFP at the predicted signal peptide cleavage site, with the exception of BLR38 that was cloned as a C-terminal fusion protein, and transiently expressed in *N. benthamiana* using *Agrobacterium*. a) Images were taken 2-3 dpi. Bars = 10 µm. n = nucleus b) Immunoblots of the fusion proteins indicating their size and stability. Most effector fusion proteins are predominantly present in monomeric form with only a single band at the expected height of the fusion protein and occasionally lower bands that may represent cleavage products. YFP-BLR20 forms an exception as besides a band corresponding to the monomer, equally intense bands are present at twice and four times the weight of YFP-BLR20 indicating that it forms (stable) dimers and multimers. Grey arrows indicate the monomeric size of the fusion proteins, black arrows indicate multimers and open arrows indicate free YFP.

**Table 3.**
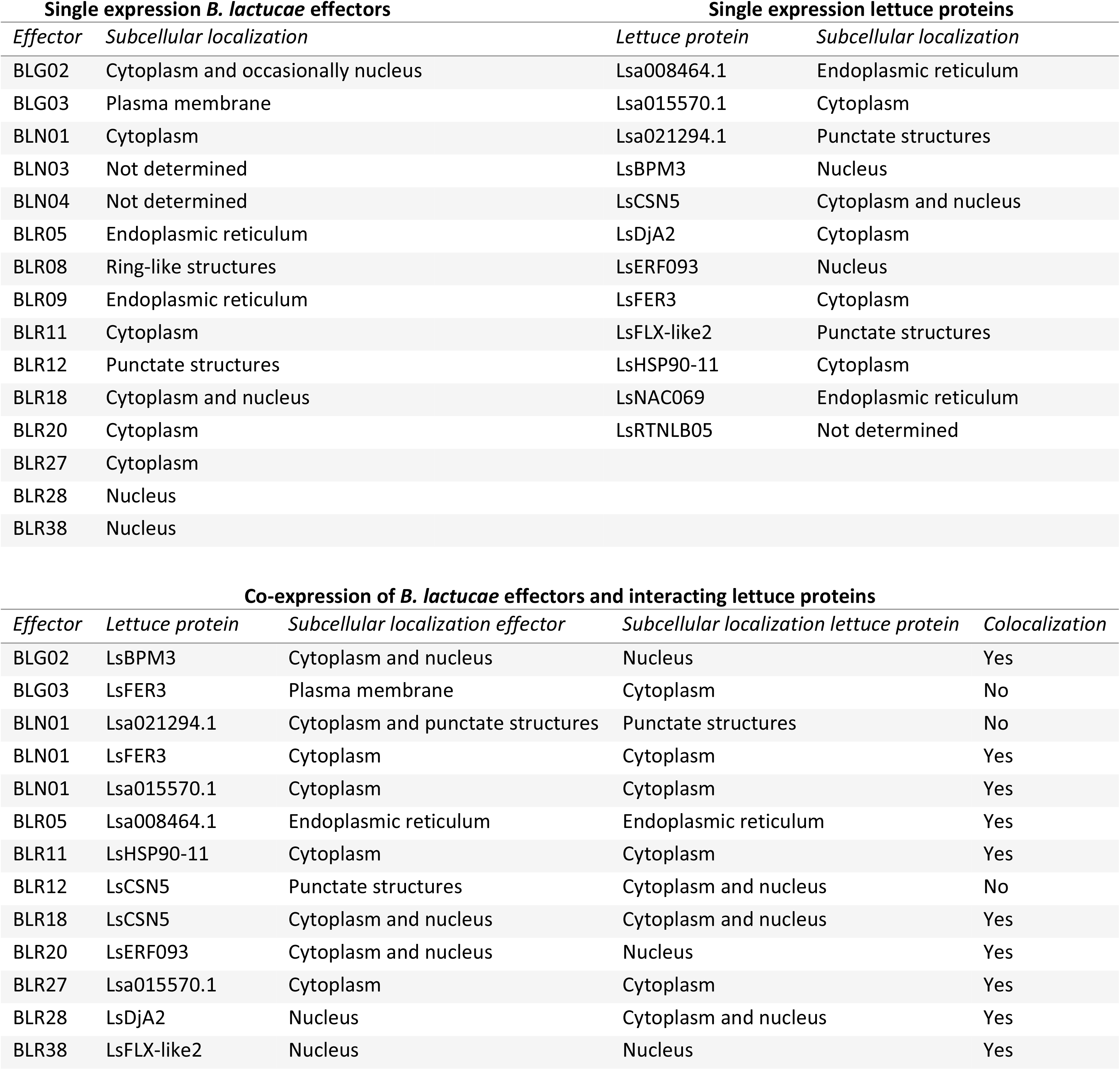
Subcellular localization of fluorescent protein fusions in *N. benthamiana*.

Subsequently, the localization of the tagged lettuce proteins was analyzed. Seven of the twelve tested proteins localized to the cytoplasm and/or nucleus (Fig 5 and Table 3). LsFLX-like2 and Lsa021294.1 localization was restricted to small, punctate structures in the cytoplasm. In line with the absence of signal peptide or transmembrane domain in LsFLX-like2 and Lsa021294.1, no ER localization was observed. The structures are thus more likely to present cytoplasmic protein complexes than components of the secretory pathway. However, for now, their origin remains undetermined. We were unable to detect C- or N-terminal LsRTNLB05 fusion proteins. LsNAC069 and Lsa008464.1 labelled the endoplasmic reticulum (Fig S2).

**Fig 5.**
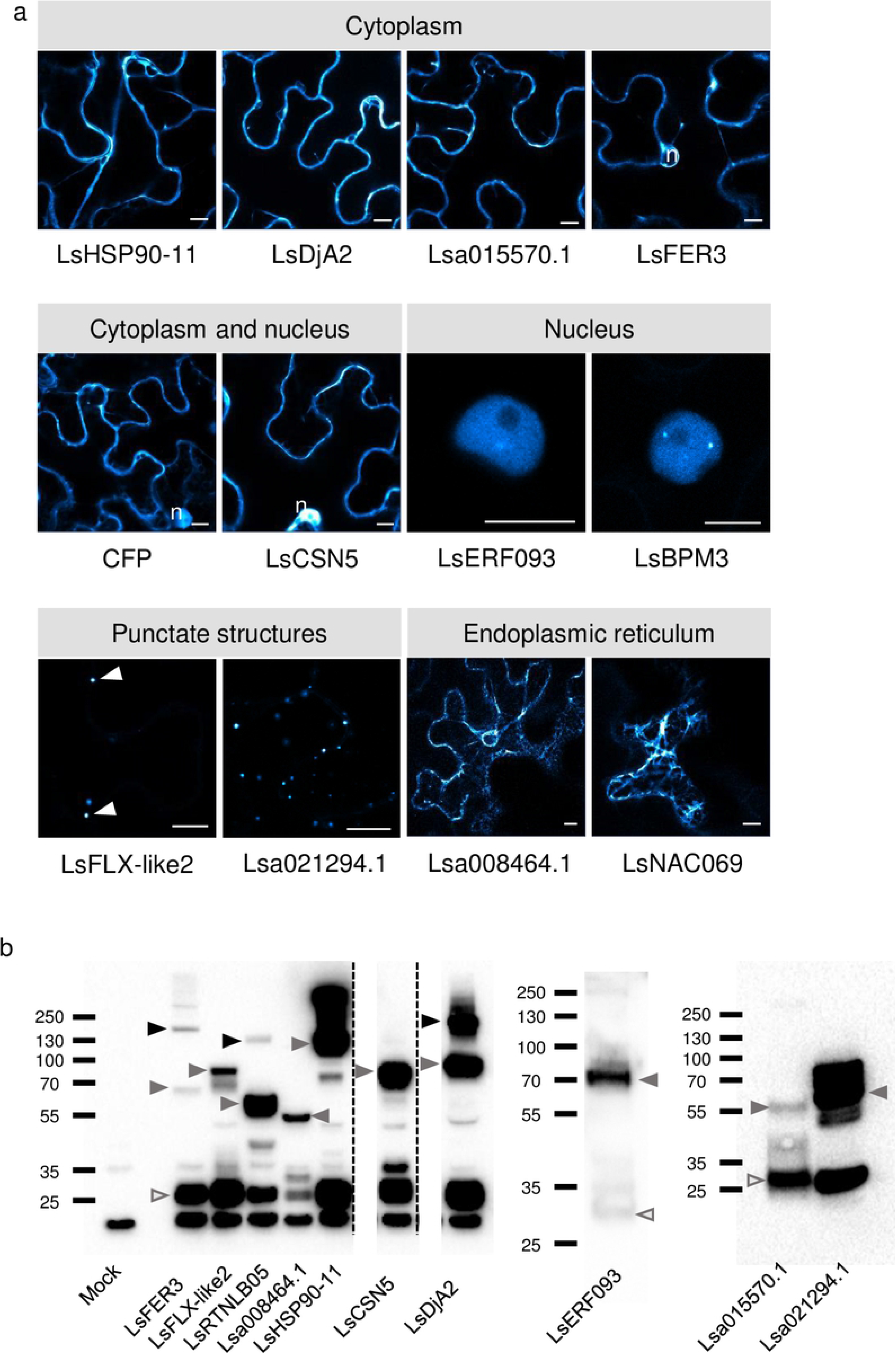
Subcellular localization and stability of effector targets in *N. benthamiana*. Lettuce proteins were fused N-terminally to CFP and transiently expressed in N. benthamiana using Agrobacterium. a) Images were taken 2-3 dpi. Bars = 10 µm. n = nucleus, arrowheads indicate the position of punctate structures. b) Immunoblots of the fusion proteins indicating their size and stability. Grey arrows indicate the monomeric size of the fusion proteins, black arrows indicate multimers and open arrows indicate free YFP.

Upon co-expression of the effectors with lettuce proteins, complete or partial colocalization was observed for 10 out of 13 combinations (Fig 6 and Table 3). As expected, the cytoplasmic localized effectors YFP-BLR11, YFP-BLR27 and YFP-BLN01 colocalized upon co-expression with their cytoplasmic targets CFP-LsHSP90-11, CFP-Lsa015570.1 and CFP-LsFER3. Interestingly, in several cases co-expression induced a relocalization, specifically, to the nucleus.

**Fig 6.**
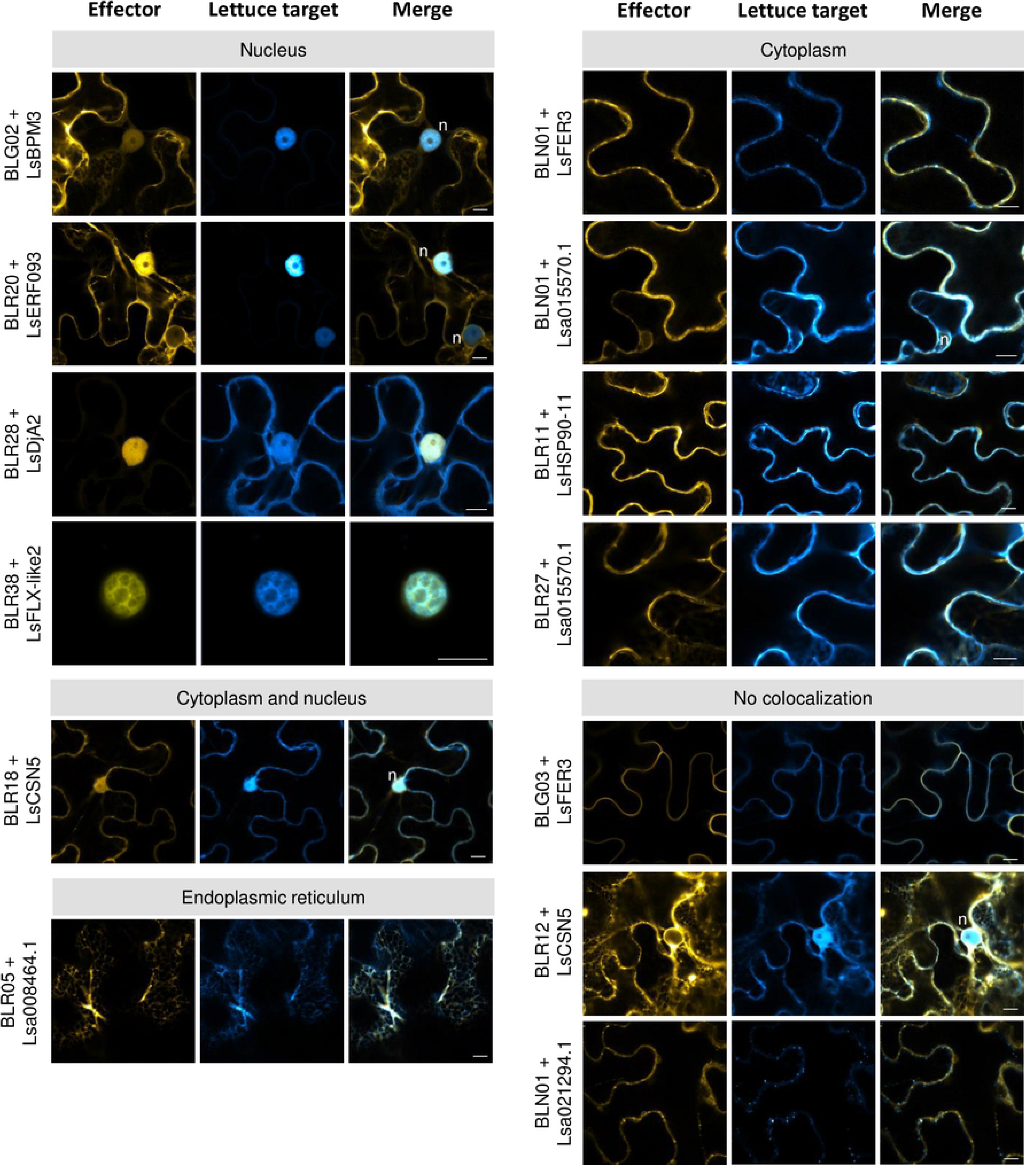
Colocalization of *B. lactucae* effectors and their targets in *N. benthamiana*. *B. lactucae* effectors and lettuce proteins were transiently co-expressed in *N. benthamiana* using *Agrobacterium*. Images were taken 2-3 dpi. Bars = 10 µm. n = nucleus.

Relocalization of effector BLG02 occurred in the presence of the nuclear localized protein CFP-LsBPM3: YFP-BLG02 was mainly cytoplasmic and, in some cells, nuclear localized, but showed a consistent nuclear and cytoplasmic (nucleocytoplasmic) signal in cells that also expressed CFP-LsBPM3 (Fig 6). The cytoplasmic effector YFP-BLR20 relocalized to the nucleus (Fig 6) upon co-expression of nuclear localized CFP-LsERF093 and the intensity of the YFP nuclear signal was dependent on CFP-LsERF093 signal intensity.

In other cases, effector expression led to target relocalization. Effector BLR38-YFP induced a relocalization of its target CFP-LsFLX-like2 from punctate cytoplasmic structures to the nucleus (Fig 6). Also, nuclear-localized effector YFP-BLR28 induced relocalization of CFP-LsDjA2 to the nucleus (Fig 6). In summary, ten of the thirteen effector-lettuce protein pairs colocalized in *N. benthamiana*. Furthermore, relocalization occurred upon expression of four effector-lettuce protein pairs providing evidence that the Y2H-identified interactions also occur *in planta*.

## Discussion

### Features of the *B. lactucae* effectors

RxLR effectors are restricted to oomycetes belonging to the clade of the Peronosporales. In each species a number of these effectors contain structurally conserved WY domains that adopt an α-helical fold and are linked to W/Y/L sequence motifs [21, 30]. The WY-domain is present in 44% of the Phytophthora RxLR effectors, but is less abundant in downy mildews: 18% of the *Plasmopara halstedii* [29] and 26 % of the *H. arabidopsidis* [21] RxLR effectors contain WY domains. Our analysis revealed that from the set of 46 *B. lactucae* effectors originating from isolate Bl:24 seven (15%) effectors have predicted WY domains. Orthologs of these seven effectors were also identified in isolate SF5 [33]. The differences in WY-domain abundance may reflect a bias in the WY-domain algorithm towards *Phytophthora* sequences; other downy mildew effectors may contain a, so far, unrecognized conserved fold.

### Yeast two-hybrid interactions between *B. lactucae* effectors and lettuce proteins

The cDNA library-based Y2H screen resulted in lettuce targets for 21 (46% of screened) effectors. These effectors interacted on average with three lettuce proteins. The vast majority (83%) of identified lettuce proteins interacted only with a single *B. lactucae* effector. Previously published screenings with *C. elegans* and human fragment libraries resulted in interactors for 37% and 31% of bait proteins respectively corresponding to an average of 2.2 and 2 interactors per bait [34, 35]. An average of 3.4 interactors per effector was obtained in screens with effectors from *G. orontii, H. arabidopsidis* and *P. syringae* against a library of ∼8000 immune-related full-length Arabidopsis proteins [8, 9]. The number of obtained interactors in our screen is thus comparable to other published Y2H screens.

Based on the identification of Arabidopsis proteins that interacted with effectors from three pathogens [9], it has been proposed that effectors converge on conserved host proteins. This proposition fits with insights into the mechanisms by which independent bacterial type III effectors converge on immune-related proteins such as MAPK proteins [36], SERK3/BAK1 [37–41] and RIN4 [42]. We were interested to determine if those proteins identified as major hubs, i.e. interacting with effectors from multiple pathogens, would also emerge in our screens with *B. lactucae* effectors.

The list of Arabidopsis hubs was dominated by TCP transcription factors [9]. Specifically, TCP13, TCP14 and TCP15 interacted with effectors originating from all three pathogens, and TCP19 and TCP21 interacted with effectors from at least one pathogen [9]. TCPs operate as transcriptional activators or repressors in plant growth and development [43, 44]. Lettuce TCP family members were also detected as interactors of *B. lactucae* effectors. However, the effector-TCP interactions weakly activated the *HIS3* reporter, were frequently observed as single colonies in Y2H screens and TCPs showed autoactivation in the Y2H system. Weßling and colleagues also classified Arabidopsis CSN5A as a hub due to its interaction with 12 *P. syringae* effectors, 11 *H. arabidopsidis* effectors and 9 *G. orontii* effectors in Y2H screens [8, 9]. In our library screens, lettuce CSN5 was identified with two *B. lactucae* effectors, but also showed weak activation of the *HIS3* reporter in the presence of an empty bait vector. Interaction between CSN5 and the GAL4 DNA binding domain was previously reported using multiple GAL4-DBD based vectors and yeast strains [45–47]. Rejecting CSN5 as false positive because of stronger reporter gene activation in the presence of *B. lactucae* effectors than empty vector is only partially defendable. Effectors may have unforeseen additive effects on reporter gene activation but these effects could be interaction independent. Thus, though some of the previously reported hubs may represent host proteins that play a prominent role in disease susceptibility, others may be false positives.

### Several *B. lactucae* effectors localize to membranes *in planta*

We examined the localization of a subset of 15 *B. lactucae* effectors and assessed their stability using immunoblot. Analysis of the YFP-BLR20 fusion protein revealed multiple bands corresponding to monomeric, dimeric and multimeric forms of this effector. It is possible that a multimer represents the biologically active form of BLR20 as multimerization of two *P. infestans* effectors, PexRD2 and PiSF3, was shown to be required for interaction with their targets, a host MAP kinase and U-Box-kinase protein respectively [21,48,49]. The multimerization of BLR20 is possibly driven by the presence of WY-domains that have been proposed to support oligomerization [21]. This was previously shown for PiSF3 by mutation of two residues facing each other across the α-helices of PiSF3 that disrupted the formation of oligomers [48].

*B. lactucae* effectors BLG03, BLN06 and BLR40, which are recognized in specific lettuce lines, localized to the plasma membrane [11, 12]. The plant membrane network was also targeted by 26% of tested *H. arabidopsidis* effectors [50] and 12% of *Plasmopara viticola* effectors [51]. Coiled-coil domains [52] and post-translational modifications such as N-myristoylation, could contribute to anchoring of effectors that lack transmembrane domains, to the plasma membrane [50,53,54]. Six effectors - BLN03, BLN04, BLR05, BLR08, BLR09 and BLR12 – were predicted to contain a single C-terminal transmembrane domain based on TMHMM analysis. Effectors BLR05, BLR08, BLR09 and BLR12 were indeed found to be associated with the endomembrane system: YFP-BLR05, YFP-BLR09 and YFP-BLR12 strongly labelled the endoplasmic reticulum, whereas BLR08 was associated with ring-like structures of varying sizes. The *B. lactucae* effectors were cloned without their signal peptide, which is required for co-translational ER targeting of transmembrane proteins by the signal recognition particle [55]. Instead membrane integration of tail-anchored proteins occurs post-translationally and is dependent on a C-terminal transmembrane domain that functions as a targeting signal. Some proteins with a moderately hydrophobic TM domain can insert unassisted in the ER, others with a highly hydrophobic TM domain require assistance. In yeast, the TM domain of tail-anchored proteins is recognized by components of the GET system including Get3 (TRC40 in mammals), delivered to the ER-resident Get1/Get2 receptor complex and inserted into the membrane [56–59]. Clearly, the ER localization of YFP-BLR05, YFP-BLR09 and YFP-BLR12 proves that these effectors associate with the membrane post-translationally. This immediately prompts the question how membrane-associated effectors are delivered to host cells. A role for extracellular vesicles in the delivery of soluble effectors has been hypothesized [60] and vesicular transport would provide a suitable vector for membrane-associated effectors as well. The C-terminal transmembrane domain of YFP-BLN03 and YFP-BLN04 may not have been sufficient for posttranslational insertion providing a possible explanation for the lack of fluorescence of these fusion proteins.

Effector BLG03 targeted the plasma membrane despite lacking a putative transmembrane domain. Post-translational modifications resulting in protein lipidation such as prenylation, N-myristoylation or palmitoylation can affect membrane-protein interactions [61]. Putative N-myristoylation sites were implicated in the membrane targeting of multiple bacterial type III effectors [54, 62]. Alternatively, phospholipid conjugation [61] or phospholipid binding may contribute to membrane association. Multiple studies have explored the role of the N-terminal RXLR motif and C-terminal residues in phospholipid binding [63]. Though the relevance of phospholipid binding for effector uptake remains unclear, perhaps a role in effector localization *in planta* could be considered.

### Co-expression of effectors and interactors induces protein relocalization to the nucleus

Studies on *H. arabidopsidis* and *P. viticola* RXLR effector localization revealed a preference for nuclear (including nucleocytoplasmic) localization (66% and 83% respectively) [50, 51]. In our research, only two effectors –BLR28 and BLR38 – were strictly nuclear localized in *N. benthamiana*. Strikingly, *B. lactucae* RXLR effector BLR38 that is recognized in *L. serriola* LS102, induced a relocalization of its target LsFLX-like2 to the nucleus. LsFLX-like2 shows homology to the Arabidopsis FLOWERING LOCUS C EXPRESSOR (FLX) protein family of which two members are involved in flowering time control in Arabidopsis [64]. The punctate structures of CFP-LsFLX-like2 resemble the structures formed by cytoplasmic FLX homodimers in Arabidopsis [65]. Heterodimers of FLX with FRIGIDA or FRIGIDA ESSENTIAL 1 were restricted to the nucleus [65], suggesting that the observed relocalization of CFP-LsFLX-like2 upon co-expression with BLR38-YFP may reflect a different interaction state. BLR28 also induced relocalization of its target, DnaJ protein LsDjA2, to the nucleus. DnaJ proteins function as co-chaperones to 70 kDa heat shock proteins to aid in protein folding, relocalization and degradation [66].

Furthermore, two predominantly cytoplasmic effectors, BLG02 and BLR20, displayed nucleocytoplasmic localization when co-expressed with their targets, LsBPM3 and LsERF093. LsERF093 contains an AP2/ERF domain that is typically found in members of the AP2/ERF transcription factor family. These proteins are known to be involved in responses to biotic and abiotic stress in Arabidopsis, tomato and rice [67, 68]. LsBPM3 contains a BTB/POZ (broad complex, tram track, bric-a-brac ⁄POX virus and zinc finger) domain that is also found in 80 Arabidopsis proteins and is known to mediate protein-protein interactions. Specifically, the BTB/POZ domain acts as a substrate-specific adaptor for Cullin-RING E3 ubiquitin Ligases (CRLs) that ubiquitinate proteins targeted for degradation by the proteasome. Recruitment of substrates may occur via a Meprin and TRAF homology (MATH) domain [69, 70]. The Arabidopsis MATH-BTB/POZ protein BPM3 localizes to the nucleus and promotes degradation of transcription factor ATHB6 that negatively regulates abscisic acid signaling [71]. Furthermore, BPMs interact with APETALA 2/ethylene-responsive element binding factor (AP2/ERF) transcription factors [72].

The relocalization events that we observed, have two important implications. First, they suggest that physical interactions between RXLR effectors and targets occur *in planta*, which may affect the activity of the target and consequently, disease development. This principle was recently demonstrated for *P. sojae* effector Avh52 that recruits a cytoplasmic host transacetylase, GmTAP1, into the nucleus. Nuclear-localized GmTAP1 is able to transacetylase histones, which enhances susceptibility of *N. benthamiana* to *P. capsici* [73]. Also, Arabidopsis protein RD19 that was discovered as interactor of *Ralstonia solanacearum* type III effector Pop2 in a Y2H screen, relocalized to the nucleus in the presence of the effector and physically associated with Pop2. The effector target, RD19, was required for the resistance protein RRS1-R mediated immune responses [74] Thus, relocalization events are often associated with physical interactions between effector and target, and effector-target interactions can affect disease susceptibility. Functional analysis of the lettuce targets is required to provide further insights into the mechanisms underlying *B. lactucae* susceptibility.

The second implication is that screening by Y2H is a useful method to identify host targets. However, the biological relevance of the observed interactions needs to be demonstrated in follow-up experiments and there is no universal rule (yet) for selecting the most promising candidates. We investigated LsFLX-like2/ BLR38 and LsDjA2/ BLR28 because they belonged to the top ten ranked interactions in the Y2H screen. In contrast, BLG02/ LsBPM3 and BLR20 / LsERF093 were selected based on literature demonstrating a role for the target candidate protein families in immune responses. It should furthermore be stressed that neither of these four targets function as hub in our Y2H screen: all of them interacted with a sole effector. Conversely, we have previously demonstrated that small networks identified in Y2H screens can be valuable i.e. LsNAC069 interacted with four - BLN04, BLR05, BLR08 and BLR09 - effectors in Y2H assays and was demonstrated to affect plant responses to both biotic and abiotic stress [32]. To conclude, our Y2H screens have uncovered multiple relevant targets whose precise function we are just beginning to unravel.

## Acknowledgements

This project (#12683) was part of the Open Technology Programme and supported by the Netherlands Organization for Scientific Research (NWO) in collaboration with Wageningen University (Wageningen, The Netherlands), Bayer, now BASF (Nunhem, The Netherlands), Enza Zaden (Enkhuizen, The Netherlands), RijkZwaan (De Lier, The Netherlands), Syngenta (Enkhuizen, The Netherlands), and Vilmorin & Cie (Lab Ménitré, France). MB was supported by the Scientific Research (NWO)/ALW Innovational Research Incentives Scheme Vidi grant 864.09.008.

## Supporting information

**S1 Table. Primers used in this study.**

**S2 Table. Prediction of importin-α-dependent nuclear localization signals**

**S3 Table. Position of transmembrane domains in effectors using TOPCONS.**

**S4 Table. Putative post-translational lipid modifications in membrane-associated effectors.**

**S1 File. Supplemental material and methods.**

**S2 File. New gene names for lettuce gene models**

**S1 Fig. Co-localization of effectors with markers in *N. benthamiana*.** A) YFP-BLR05 and YFP-BLR09 co-localize with an CFP-tagged ER lumenal marker. B) YFP-BLG03 localizes to the plant plasma membrane. Leaf sections were incubated in propidium iodide (PI) to stain the cell wall. Bars = 10 µm.

**S2 Fig. LsNAC069 and Lsa008464.1 co-localize with an ER lumenal marker in *N. benthamiana*.** Bars = 10 µm.

**S3 Fig. Phylogenetic tree of lettuce BTB/POZ domain containing proteins.** The Y2H identified target, LsBPM3, forms a distinct clade with four other lettuce BTB/POZ domain containing proteins and six Arabidopsis BPM proteins. AtBPM1-AtBPM6 contain in addition to a BTB/POZ domain also a MATH domain. Protein sequences were aligned using Clustal Omega, a Neighbor-Joining phylogenetic tree was constructed using MEGA7 and the tree was visualized using iTOL software.

**S4 Fig. Phylogenetic tree of lettuce and Arabidopsis FLX(-like) proteins.** Protein sequences were aligned using Clustal Omega, a Neighbor-Joining phylogenetic tree was constructed using MEGA7 and the tree was visualized using iTOL software.

